# CTCF mediated genome architecture regulates the dosage of mitotically stable mono-allelic expression of autosomal genes

**DOI:** 10.1101/178749

**Authors:** Keerthivasan Raanin Chandradoss, Kuljeet Singh Sandhu

## Abstract

Mammalian genomes exhibit widespread mono-allelic expression of autosomal genes. However, the mechanistic insight that allows specific expression of one allele remains enigmatic. Here, we present evidence that the linear and the three dimensional architectures of the genome ascribe the appropriate framework that guides the mono-allelic expression of genes. We show that: 1) mono-allelically expressed genes are assorted into genomic domains that are insulated from domains of bi-allelically expressed genes through CTCF mediated chromatin loops; 2) evolutionary and cell-type specific gain and loss of mono-allelic expression coincide respectively with the gain and loss of chromatin insulator sites; 3) dosage of mono‐ allelically expressed genes is more sensitive to loss of chromatin insulationn associated with CTCF depletion as compared to bi-allelically expressed genes; 4) distinct susceptibility of mono‐ and bi-allelically expressed genes to CTCF depletion can be attributed to distinct functional roles of CTCF around these genes. Altogether, our observations highlight a general topological framework for the mono-allelic expression of genes, wherein the alleles are insulated from the spatial interference of chromatin and transcriptional states from neighbouring bi-allelic domains via CTCF mediated chromatin loops. The study also suggests that the three-dimensional genome organization might have evolved under the constraint to mitigate the fluctuations in the dosage of mono-allelically expressed genes, which otherwise are dosage sensitive.

## Introduction

Though both copies of genes on autosomes have potential to be expressed, some genes escape the transcriptional activation of one of the alleles. These genes are known as mono-allelically expressed (MAE) genes. MAE genes can be genomically imprinted, i.e., the expressions of alleles are parentally fixed, or it can be random, i.e., any of the alleles can be expressed. A subset of random MAE genes has been shown to be mitotically stable in a clonal cell population, possibly through heritable epigenetic modifications of alleles[1]. Intriguingly, random MAE genes have profound functional and evolutionary implications such as contributing to cellular diversity of immunoglobulins[2], interleukins [3, 4] and T-cell receptors[5]; ascribing choice of olfactory receptors in neurons[6] and increasing the evolvability of a locus[7]. Despite widespread presence of MAE genes in mammalian genomes, the underlying mechanisms that regulate allele-specific transcription remain poorly understood. Feedback mechanism in which expression of one allele induces the repression of other allele seems an interesting hypothesis. It has been shown that expression of a transgene odorant receptor in neurons leads to lack of endogenous expression of odorant receptor alleles, which is contrasting when compared to biallelically expressed genes which exhibit co-expression with the transgenes[8]. Though appealing, the mechanistic details of this phenomenon remain enigmatic. Directional switching of an upstream promoter has been shown to stochastically regulate the activity of downstream promoter of NK receptor gene in an allele‐specific manner[9]. Epigenetic mechanisms like differential methylation of alleles[10, 11], non-coding RNA mediated repression[12, 13], distinct spatial localizations of alleles[14, 15] etc. have also been proposed and exemplified for certain MAE loci. While these mechanisms are supported through experimental evidence, none of these could be generalized for most of the mono-allelic transcription of the genome. Moreover, any of the gene regulatory mechanisms implicated in regulating the two alleles distinctly would need a prerequisite of recognizing the MAE genes from the neighbouring BAE genes and one way this can be achieved is by insulating the MAE genes from BAE genes on either or both allelic loci. We, therefore, tested the hypothesis whether or not CTCF mediated insulation of chromatin domains implicate in guiding mono-allelic expression in the genome. Testing such a hypothesis at genome scale is constrained by the genomic coverage of MAE and BAE genes, which has been limited due to limited availability of allelic variants in the coding regions of the genes. However, in a recent pioneer study it has been demonstrated that MAE and BAE genes can be reliably identified on the basis of simultaneous occurrence of transcription elongation mark H3K36me3 and repressive mark H3K27me3 in the gene-bodies[16]. This led to higher genomic coverage for MAE and BAE genes, which allows for unbiased genome-wide analyses. Our systematic analyses of experimentally identified and inferred MAE/BAE genes revealed a possible role of chromatin topology in ascribing appropriate framework for the mono-allelic expression of genes and suggested that the CTCF mediated dosage control of MAE genes can serve as a potent constraint shaping the evolution of linear and 3D genome organization.

## Results

We obtained experimentally identified as well as inferred mitotically stable MAE and BAE genes in human lymphoblastoid cell-line (hLCL), mouse lymphoblastoid cell-line (mLCL) and mouse embryonic stem cells (mESC) (Table 1). To ensure that our analysis was not impacted by the already known properties, like clustering of imprinted genes[17], we removed the known imprinted loci from the present analysis. Possible regulatory differences of MAE and BAE genes prompted the hypothesis that the mono‐ and bi-allelic expression might be a domain property, of i.e., MAE genes might tend to segregate from BAE genes in a manner similar to imprinted genes. To test this, we calculated the observed frequencies of MAE-MAE, BAE-BAE and mixed (MAE-BAE/BAE-MAE) gene-pairs across the human and mouse genomes and compared with that of expected frequencies (Materials and Methods). As shown in the Figure 1a, MAE-MAE and BAE-BAE pairs were clearly over‐ represented as compared to mixed pairs in human and mouse cell-types, suggesting that MAE and BAE genes are assorted from each other in the genome. To further confirm this observation, we calculated the density of MAE and BAE genes across chromosomes and identified regions enriched with MAE, BAE or both type of genes (Materials & Methods). We showed that the gene clusters with either MAE or BAE genes were the majority as compared to ones with both, highlighting the preferred segregation of MAE and BAE domains (Figure 1b, S1a). Through randomizing the MAE and BAE labels of genes, we reported that the observed clustering of MAE and BAE genes was highly non-random (p-value=2.2e-16, Chi-squared test, Materials & Methods). Further, to test if active and inactive alleles of MAE genes were also clustered separately, we obtained paternally active (PMAE) and maternally active MAE (MMAE) genes in hLCL (Table 1). Analysis of PMAE and MMAE suggested that active and inactive alleles did not exhibit random mixing and were mostly clustered separately (Figure S1b). These observations suggested that the linear gene order might have evolved under the evolutionary constraint to segregate MAE genes from BAE genes.

**Figure 1.**
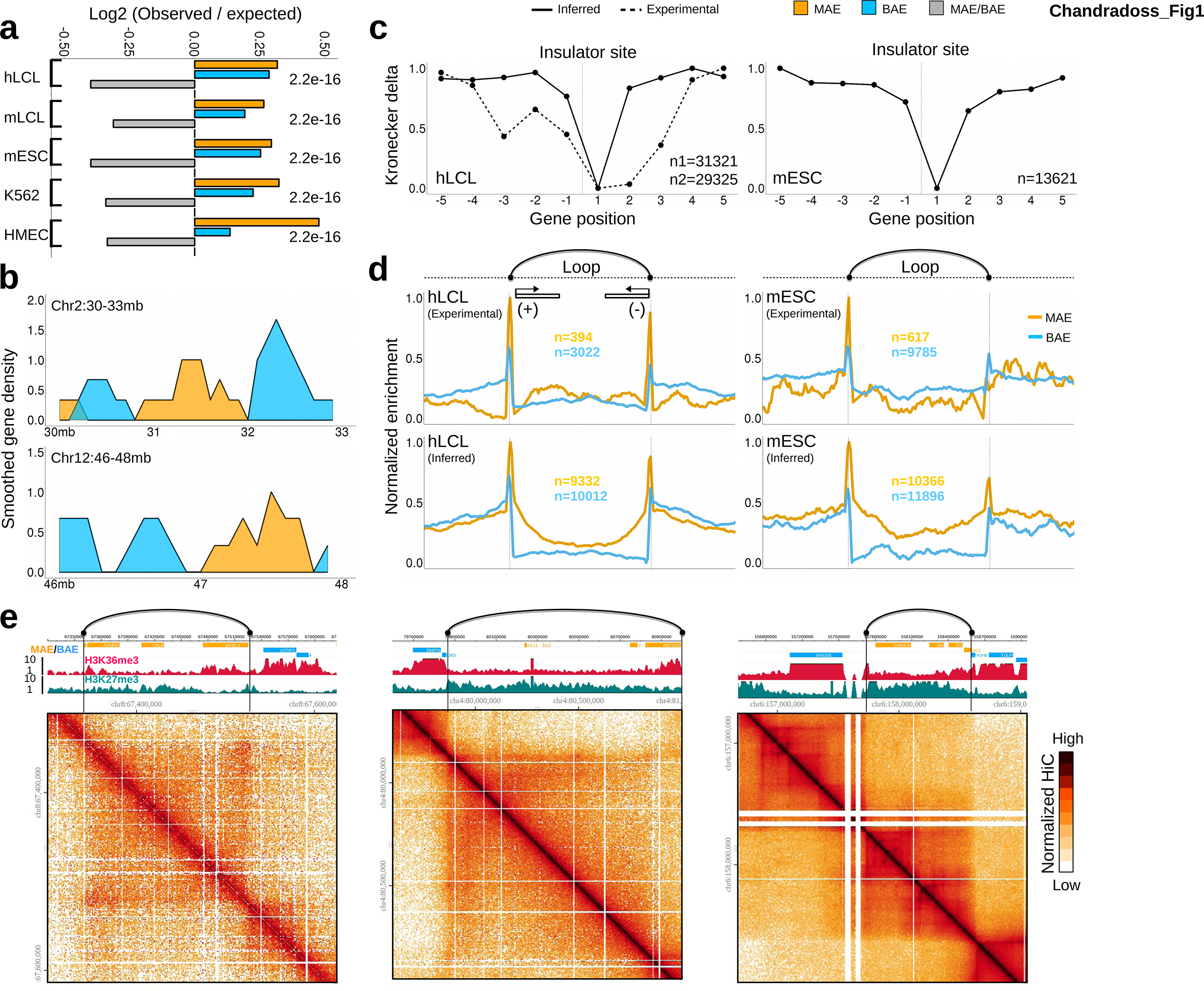
Genomic assortment of MAE and BAE genes. (a) Observed-to-expected ratio of MAE-MAE, BAE-BAE and mixed gene pairs for different datasets. P-values are calculated using Chi-squared test for observed and expected counts. (b) Examples illustrating the linear compartmentalization of MAE and BAE genes. Chromosome coordinates are given for hg19 assembly. (c) Scaled average aggregation plot of Kronecker delta function over five consecutive genes upstream and downstream to insulator CTCF sites. (d) Normalized enrichment of MAE and BAE genes inside and around chromatin loops mediated by insulator CTCF sites. (e) Example snapshots from WashU Epigenome Browser showing MAE genes (orange bars), BAE genes (blue bars), H3K36me3 (red), H3K27me3 (green), CTCF ChIA-PET loops (black arcs) and TAD domains (heatmaps, 25/50 kb resolution). Shown regions are chr8:67.3-67.625 Mb, chr4:79.6-81 Mb and chr6:156.67-159.05 Mb in hLCL (hg19).

**Table 1.** Overall statistics of MAE and BAE genes taken for the analysis.

| Method | Species | Cell-line | MAE | BAE | Source |
| --- | --- | --- | --- | --- | --- |
| Experimentally identified | Human | Lymphoblastoid | 399 | 3045 | Gimelbrant et al [1] |
|  | Mouse | Pre-B | 294 | 1101 | Zwemer et al [35] |
|  |  | ESC | 629 | 9827 | Gendrel et al [42] |
| Inferred | Human | Lymphoblastoid | 9469 | 10085 | Nag et al[16] |
|  |  | HMEC | 6529 | 12880 | Nag et al [16] |
|  |  | K562 | 8574 | 10455 | Nag et al [16] |
|  | Mouse | ESC | 10427 | 11955 | Nag et al[43] |
|  |  | Lymphoblastoid | 8967 | 11647 | Nag et al[43] |
| Experimentally identified | Human | Lymphoblastoid | **MMAE** | **PMAE** |  |
|  |  |  | 484 | 422 | Rozowsky et al [44] |

Given that the MAE genes taken in the analysis were mitotically stable, we further hypothesized that the domains of MAE and BAE genes might be epigenetically insulated from each other. CTCF protein is presently the most popular candidate that serves as insulator between epigenetically distinct domains. We, therefore, tested the presence of CTCF binding at boundary of MAE and BAE domains. We obtained the CTCF binding sites that were associated with the insulator state in chromHMM annotations of hLCL and mESC genomes (Materials and Methods). We calculated Kronecker delta function (**δ(x_i_, x_i+1_)**, where **x_i_** is the allelic status of the gene i) for consecutive genes around CTCF insulator sites. The function took the value 1 if a MAE gene was followed by a MAE gene or a BAE gene was followed by a BAE gene, otherwise the value remained zero. It was clear from the Figure 1c that the allelic status of the gene changed after encountering a CTCF insulator site. This was also exemplified through numerous examples shown in the other panels of Figure 1. Moreover, PMAE and MMAE genes also exhibited similar pattern as shown in figure S1c. We, therefore, conclude that the mono‐ or bi-allelic expression is the property of chromosomal domains, insulated by CTCF insulator sites, rather than individual genes.

CTCF orchestrates the genome in defined topological domains that are intervened by inter-domain or gap regions. To obtain further insight, we mapped the locations of MAE and BAE genes within gap-loop-gap architecture obtained from CTCF ChIA-PET data of hLCL and mESC. We observed that the MAE genes were preferably located inside the chromatin loop between insulator CTCF sites, while BAE genes exhibited lesser such preference (Figure 1d). To test if our observation was not merely a consequence of different expression levels of MAE and BAE genes, we sampled MAE and BAE genes of similar expression levels and reassessed their association with the insulator loops. We confirmed that the MAE genes were preferably associated with the insulator loops as compared to BAE genes of similar expression levels (Figure S2). This hinted that the insulator CTCF sites proximal to MAE genes might have been implicated and evolutionarily selected to maintain the mono-allelic expression. We, therefore, tested if the dynamics of allelic expression correlated with the gain and loss of insulator function of CTCF binding site in the proximity. Towards this, we compared the MAE and BAE genes in hLCL and K562 cell-lines. We first showed that the constitutive MAE genes (genes that maintained their mono-allelic status consistently in both cell-lines) exhibited consistently greater enrichment inside the chromatin loops mediated by insulator CTCF sites as compared to constitutive BAE genes in both the cell-lines (Figure 2a). Interestingly the genes that were MAE in hLCL, but had bi-allelic expression in K562 cell-line, had significant enrichment inside the chromatin loop in hLCL, but not in K562 cell-line (Figure 2b). This pattern reversed for the genes that were mono-allelic in K562 but had bi-allelic expression in hLCL (Figure 2b). The observed gain and loss of enrichment of MAE genes inside the chromatin loops mediated by CTCF insulator can be explained by the difference in CTCF mediated loops (32% of the cases) and the relative chromatin context of CTCF binding sites (68% of cases) in the two cell-lines. This was illustrated through examples: 1) CTCF mediated loop around a locus that was MAE in hLCL, was not observed in K562 where the gene was expressed bi-allelically. By plotting RNAPII ChIA-PET data, we clearly observed lack of insulation and gain of abundant enhancer-promoter and promoter-promoter interactions with the neighbouring regions in K562 cell-line (Figure 2c). 2) CTCF mediated loop remained intact in both the cell-lines, but the chromatin context of CTCF binding differed. In the cell-line where the gene was expressed bi-allelically (hLCL in this case), the promoter of the gene gained additional CTCF binding sites, which were engaged with the other CTCF and non-CTCF sites associated with enhancer chromatin states (Figure 2d). Indeed, the example shown in the figure 2d suggested that the CTCF sites proximal to ITPKB gene-promoter interacted with up/downstream enhancers and the 3’ termination site of the gene highlighting the transcriptionally active gene complex widely claimed elsewhere[18-21]. It was interesting to note that the CTCF binding sites that enclosed MAE genes into insulator loops in one cell-line, can function as enhancer/terminator-linker for the bi-allelic expression of the gene in the other cell‐line. Indeed, BAE genes were significantly associated with the enhancer-linking CTCF sites, as compared to MAE genes (Figure S3). The observed correlation between mono-allelic status of genes and their localization near CTCF insulator sites strongly supported the role of CTCF insulators in maintaining mono-allelic expression of genes. We also confirmed the above observations by comparing hLCL with the HMEC cell-line (Figure S4).

**Figure 2.**
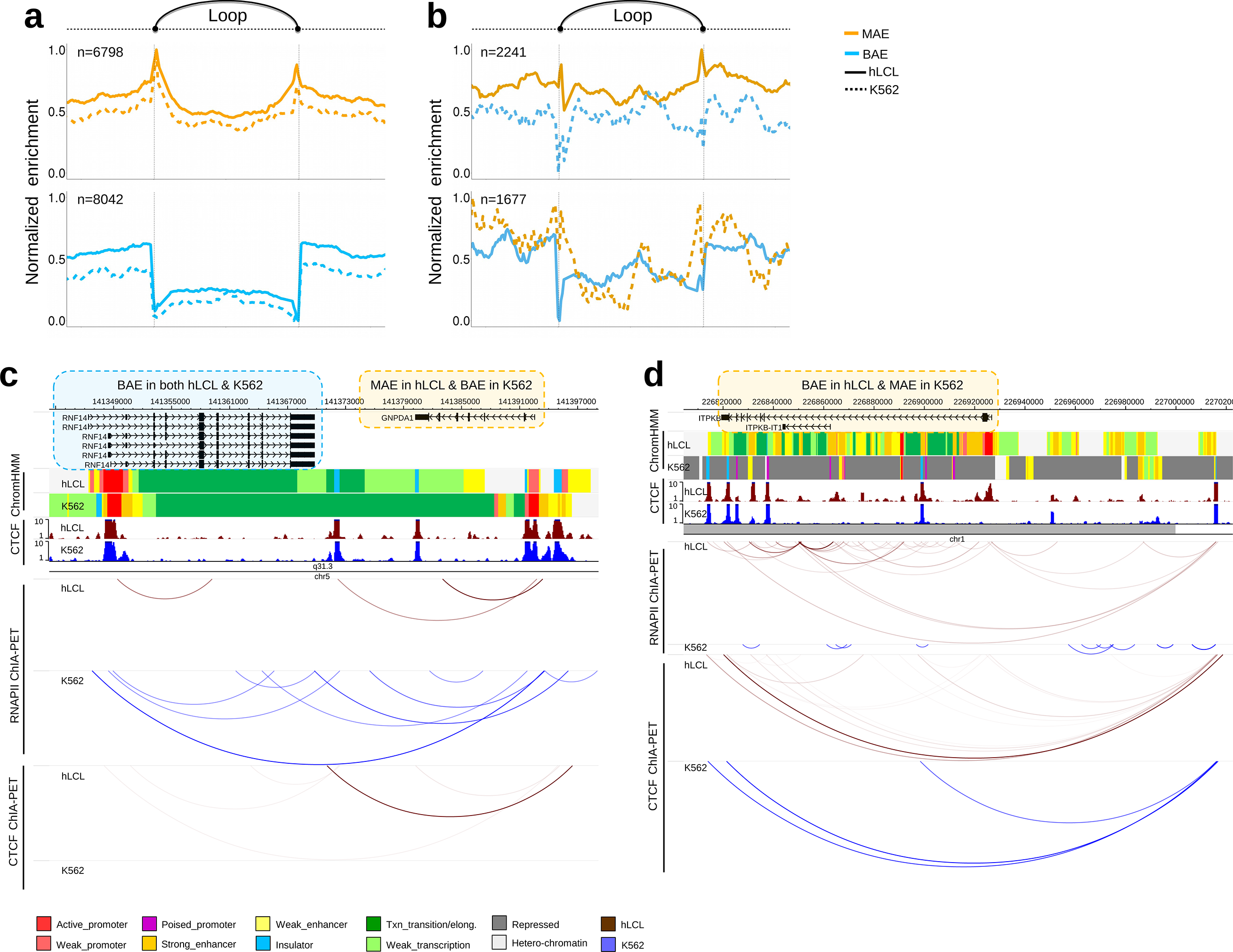
Cell-type specific association between allelic status of genes and chromatin architecture mediated by CTCF insulator. (a) Enrichment of MAE and BAE genes that maintained their allelic status in hLCL and K562 cell-lines, inside and around insulator loops (b) Enrichment of MAE and BAE genes that switched their allelic status in hLCL and K562 cell-lines, inside and around insulator loops. (c) Examples illustrating switch in allelic status of gene between two cell-lines. Shown are the RefSeq genes, chromHMM, CTCF ChIP-Seq, RNAPII ChIA-PET and CTCF ChIA-PET tracks for hLCL and K562 cell-lines. In the left panel, GNPDA1 gene was MAE in hLCL and BAE in K562 cell-line. In the right panel, ITPKB gene was BAE in hLCL and MAE in K562 cell-line.

Further, to assess whether or not gain of CTCF insulator sites near MAE genes was evolutionarily selected, we compared the MAE and BAE genes of mLCL with that of hLCL. The genes that maintained their mono-allelic expression in human and mouse LCLs were consistently associated with the CTCF insulator sites in the proximity (Figure 3a). However, the evolutionary loss/gain of mono-allelic expression coincided with the loss/gain of insulator sequence in the proximity (Figure 3b-c). These observations highlighted two things: 1) Greater association of MAE genes with CTCF insulator sites appears to be evolutionarily conserved property of mammalian genomes. 2) Gain of genomic proximity to some insulator sites might have been evolutionarily selected to maintain mono-allelic expression of genes. Altogether, observations through figure 2 and 3 suggested that the genetic and the epigenetic association with the CTCF insulator sites might serve as a prerequisite for the mono‐ allelic expression of the genes. Since the loss of CTCF insulator function coincided with the gain of bi-allelic expression, it could also be inferred that CTCF insulator function was associated with the repressed alleles of MAE genes.

**Figure 3.**
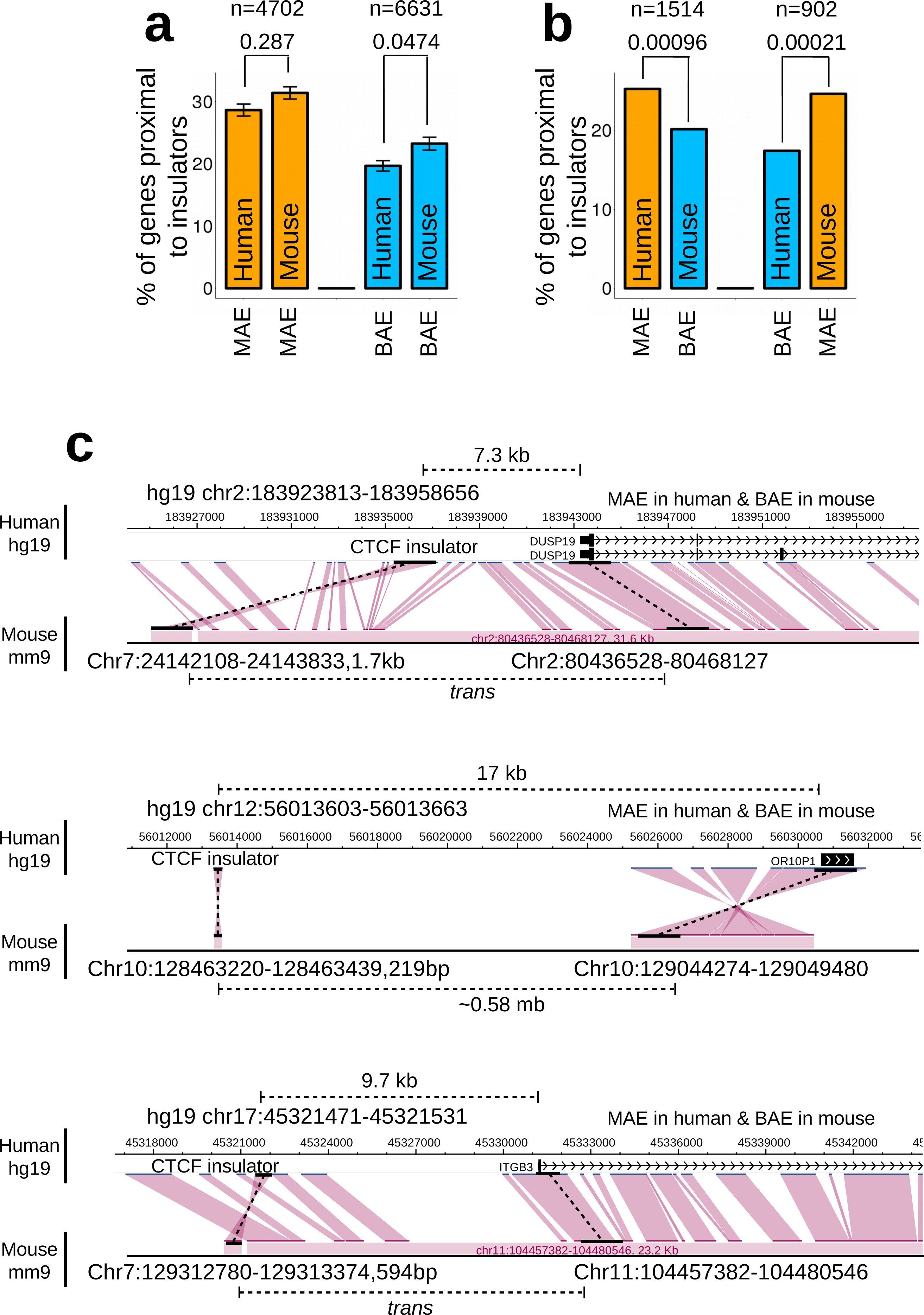
Genetic association between allelic status of genes and their proximity toinsulator CTCF sites. (a) Proportion of MAE and BAE genes that maintained their allelic status in human and mouse lymphoblastoid cell-lines and were having at-least one insulator CTCF site within 20kb upstream to TSS. (b) Proportion of MAE and BAE genes that switched their allelic status in human and mouse lymphoblastoid cell‐ lines and were having at-least one insulator CTCF site within 20kb upstream to TSS. P-values were calculated using Chi-squared test. (c) Examples illustrating association between gain/loss of genomic proximity to insulator CTCF sites and the allelic status of the gene in human and mouse genomes. Snap-shots were obtained from WashU Epigenome Browser.

With the recent availability of genome-wide CTCF depletion datasets, it is now possible to explore if expression and insulation of certain predefined subset of genes is affected by the loss of CTCF function. We obtained CTCF depletion datasets for mLCL and mESC (Materials and Methods). We first showed that the CTCF depletion disrupted the insulation of MAE and BAE domains significantly (Figure 4a, d). Importantly, the disruption was more striking for MAE genes. We further showed that the dosage of MAE genes was strikingly more sensitive to CTCF depletion as compared to that of BAE genes (Figure 4b-d). The higher sensitivity of MAE genes was also observed for depletion of Cohesin loader Nipbl (Figure S5). It was noticeable that the MAE genes that were up-regulated after CTCF depletion outnumbered the ones that were down-regulated when compared to BAE genes, again suggesting that the repressed allele was likely to be associated with insulator mediated chromatin loops (Figure S6). Further, the up‐ and down-regulation of MAE genes coincided with the coherent transcriptional states in the neighbouring domains. The chromatin loops with up-regulated MAE genes were flanked by active genes as compared to the ones flanking the loops with down-regulated MAE genes. Importantly, the promoters of up-regulated MAE genes exhibited increased interaction frequencies with the enhancers and enhancer-like promoters from the neighbouring domains upon CTCF depletion. In contrast, promoters of down‐ regulated genes exhibited increased interactions with the repressed promoters from the neighbouring domains upon CTCF depletion. These observations suggested that the lack of insulation between neighbouring chromatin domains might have impacted the allelic status of MAE genes through altered chromatin interactions (Figure 5a-b).

**Figure 4.**
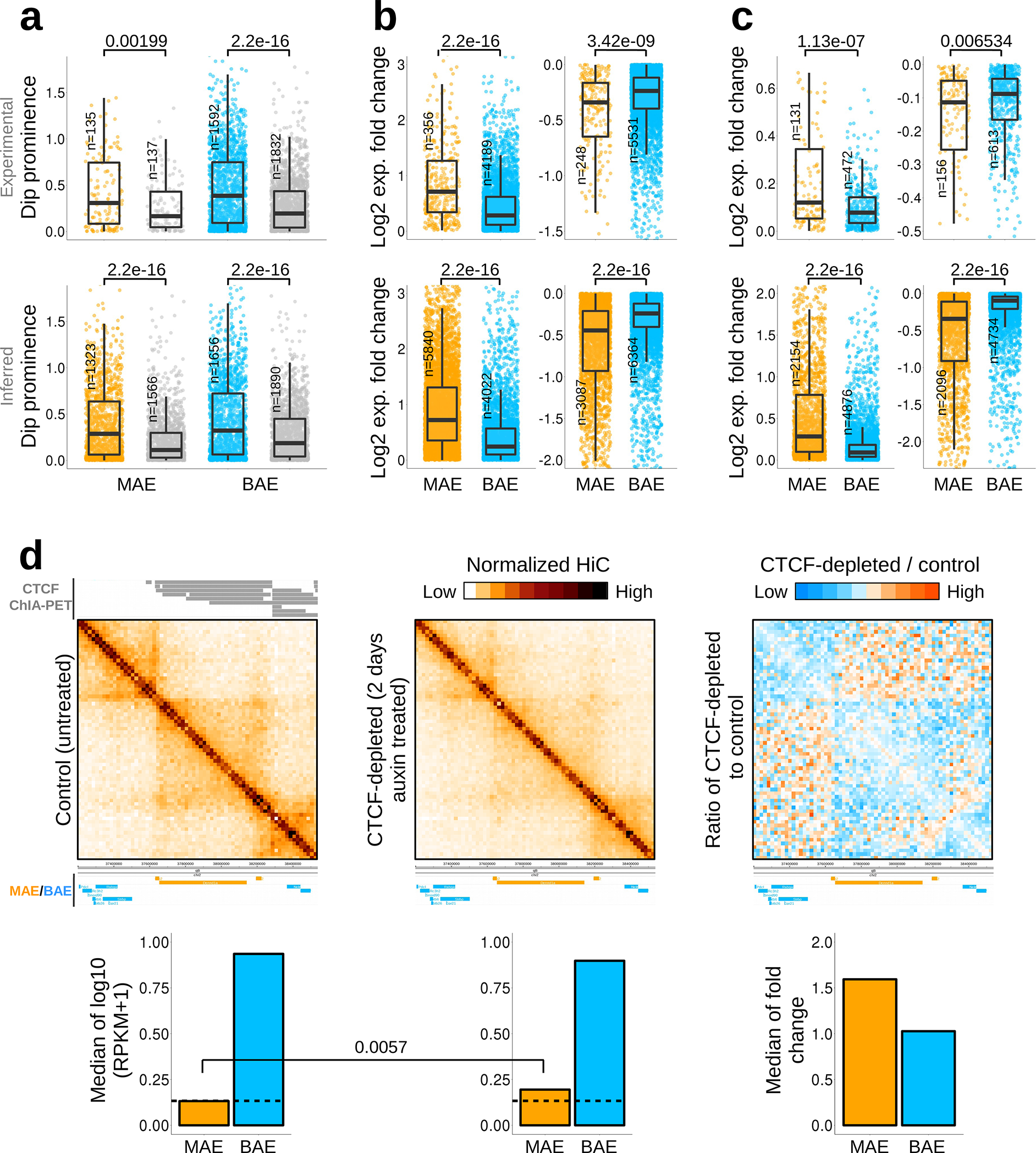
Impact of CTCF depletion on MAE and BAE genes. (a) Boxplots of dip prominence (insulation) of MAE and BAE genes in the control and CTCF depleted mESC. (b-c) Log2 fold change of expression of MAE and BAE genes after CTCF depletion in (b) mESC, and (c) mLCL. P-values were calculated by Mann-Whitney U test. (d) An example illustrating chromatin insulation of MAE genes in mESC and lack thereof after CTCF depletion. Shown are MAE, BAE genes; CTCF ChIA-PET loops; the heatmaps for HiC data before CTCF depletion, after CTCF depletion and the ratio of the two; gene expression levels of MAE/BAE genes in the locus before CTCF depletion, after CTCF depletion and the ratio of the two.

**Figure 5.**
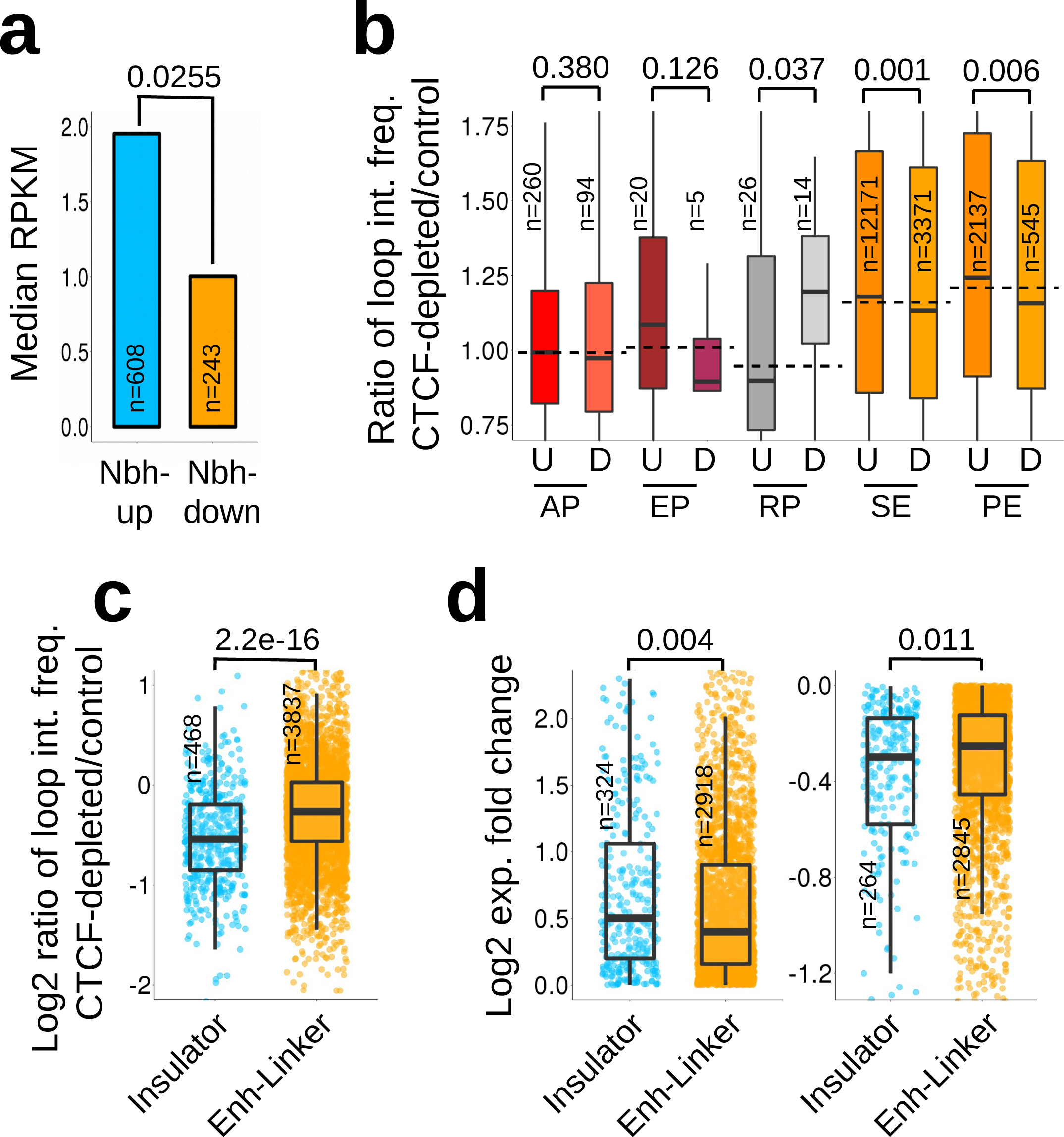
Mechanisms attributing to susceptibility of MAE genes to CTCF depletion (a) Median expression of genes, which were located in the neighbouring chromatin domains of up-regulated and down-regulated MAE genes. P-value was calculated using Mann Whitney U test (b) Fold change in frequencies of interactions between promoters of up/down-regulated MAE genes and different kinds of promoters and enhancers in the neighbouring loops after CTCF depletion. The dotted lines represent the genome-wide median change for each chromatin state. P-values were calculated using Mann-Whitney U test. (c) Fold change in interaction frequencies of all genes enclosed within chromatin loops mediated by insulator CTCF sites and the genes with their promoters linked to enhancers via enhancer-linking CTCF sites. Mutually exclusive sets are compared (d) Fold change in expression of genes associated with insulator loops and enhancer linking loops before after CTCF depletion in mESC.

It was not entirely clear why the dosage of MAE genes was more sensitive to CTCF depletion. We suspected that the insulator function of CTCF might be more susceptible to CTCF depletion as compared to enhancer-linking. To test this, we compared the interaction frequencies of CTCF mediated enhancer-promoter loops with that of insulator anchored loops before and after CTCF depletion in mESC. We observed relatively lesser alteration in interaction frequencies of enhancer-linker sites after CTCF depletion as compared to the loops anchored to insulator sites (Figure 5c), suggesting that the minimal amount of CTCF was sufficient to link enhancer to their cognate promoter. Accordingly, the expression of genes associated with insulator loops were more susceptible to CTCF depletion as compared to the ones associated with enhancer-linking loops (Figure 5d). Since MAE genes were flanked by insulator sites, while BAE genes were enriched near enhancer-linking CTCF sites, it can be concluded that distinct functional roles of CTCF around MAE and BAE genes might explain distinct susceptibility of MAE and BAE genes to CTCF depletion. One way, such an effect can take place is through deposition of multiple transcription factors around enhancer-linking CTCF sites, which can stabilize CTCF binding through cooperative binding as opposed to that around insulator site. Though hypothetical, the above extrapolation is based on the fact that the enhancer sites are known to accommodate clusters of TF binding sites[22]. It can also be argued that MAE genes are more likely to be transcriptionally perturbed as compared to BAE genes, because only one allele needs to be dysregulated. We attempted to address this by assessing the proportion of MAE and BAE genes that were dysregulated across different publicly available perturbation experiments in mESCs. The mean proportion of MAE genes that got dysregulated across different perturbations did not exceed significantly with that of BAE genes, suggesting that the sensitivity of MAE genes to CTCF/Cohesin depletion is independent of presumed sensitivity of MAE genes to any transcriptional perturbation (Figure S7).

## Discussion

CTCF mediated chromatin folding is known to implicate in maintaining allele-specific transcriptional states of H19-Igf2 imprinted locus. Loss of CTCF binding at H19-ICR leads to loss of maternal repression of Igf2, which otherwise is insulated in a repressive loop mediated by CTCF[23]. Our observations suggested certain level of generality of insulation of inactive allele during mono-allelic expression. Indeed, we tested whether CTCF mediated chromatin loops were associated with the repressed alleles of MAE genes in a manner that was analogous to allele-specific repression of Igf2. Towards this, we obtained the haplotype resolved HiC data from Rao et al [24]. We first showed that the CTCF barrier loops associated with MAE genes exhibited greater variation between homologous chromosomes as compared to the ones associated with BAE genes, suggesting allele-specificity of chromatin conformation associated with MAE genes (Figure S8a). We inferred the maternally and paternally expressed alleles by making use of RNAPII ChIA-PET data (Materials and methods). By comparing the CTCF associated allele-specific chromatin interactions around maternally and paternally expressed genes, we showed that the inactive allele had relatively higher interaction frequency of CTCF mediated chromatin loop as compared to active allele, reconciling our proposal that the CTCF associated insulation was mostly associated with the inactive alleles (Figure S8b). However, due to subtle statistical difference seen in the analysis, we do not entirely deny the possibility that the up-regulation of MAE genes upon CTCF depletion might not necessarily relate to repressive allele and instead the active allele might get further up-regulated. Indeed, despite the fact that CTCF binding at H19-ICR is critical to establish and maintain proper H19-Igf2 imprinting, depletion of CTCF protein itself does not associate with the loss of mono-allelic expression of Igf2 and instead causes increased expression of the active allele itself, possibly via altering CTCF binding at other nearby sites[25]. This suggested that the minimal amount of CTCF is sufficient to maintain the imprinting at H19-Igf2 locus and that the dosage of imprinted gene can be susceptible to CTCF depletion in a non-allelic manner. We, therefore, largely restrict our claim to dosage sensitivity of MAE genes to the loss of CTCF mediated insulation around the locus.

Despite being relatively fewer in numbers, explanation to down-regulated genes was needed. It can be interpreted that the down-regulation was that of active allele. There can be following possibilities: 1) Association with the CTCF mediated chromatin loops, though statistically significant, might not always be related to repressed allele and at certain MAE loci, it might associate with active allele instead. Indeed, as shown in the figure 5b, the down-regulated MAE genes had gained interactions with the repressed promoters in the neighbouring loops after CTCF depletion, which was in sharp contrast to loops with up-regulated MAE genes, suggesting that the increased spatial interference with the neighbouring repressive domains might cause down-regulation of active alleles of MAE genes; 2) Gene regulatory network downstream to dysregulated transcription factors can also be a possibility. Regardless of the fact whether the MAE genes were up-regulated or down-regulated or whether the changes were allelic or non-allelic, significantly greater dosage sensitivity of MAE genes to CTCF depletion as compared to BAE genes is a novel and non-trivial observation, which has implication in understanding the constraints that shape the evolution of genome architecture. It has been hypothesized earlier that the MAE genes are likely to be dosage sensitive[18]. We tested this hypothesis using dosage sensitivity and copy number variation data in human. MAE genes had significantly greater overlap with the dosage sensitive genes obtained from ClinGen database (Materials & Methods, Figure S9a). Accordingly, BAE genes had greater overlap with the regions exhibiting common copy number variation in human (Figure S9b), suggesting their general insensitivity to dosage. Therefore, it can be inferred that evolutionary selection of appropriate dosage of MAE genes might have constrained the CTCF mediated chromatin insulation architecture of the genome.

Nora et al recently reported that the CTCF depletion impacts the transcriptional states but the chromatin states, as measured through H3K27me3, remains largely unaltered [26]. We also confirmed this in the context of the present study (Figure S10). Though the insulation and expression levels were significantly altered, H3K27me3 levels showed only subtle changes after CTCF depletion. Accordingly we suggest that the loss of insulation of TAD boundaries might have introduced spatial interference of opposite chromatin and transcriptional states in the neighbourhood in the form of altered enhancer-promoter and promoter-promoter interactions across TAD boundaries, which might have impacted the expression of MAE genes. The BAE genes, on the other hand, were mostly flanked by the CTCF sites involved in enhancer-linking function, which surprisingly were relatively robust against CTCF depletion as compared to insulation. As a result, BAE genes exhibited comparatively lesser transcriptional dysregulation upon CTCF depletion.

The major criticism of the work could be the use of inferred MAE and BAE genes. It can be argued that the simultaneous presence of H3K27me3 and H3K36me3 marks might not necessarily mark the mono-allelic genes and rather represent the instances of cellular heterogeneity of these marks, particularly in mESCs that exhibit heterogenous expression of developmental regulators mostly due to serum components of the culture[27, 28]. In this regard, we emphasize the following points: 1) Nag et al has shown experimental validation through RNA-seq and AST-seq of their proposed inference method[16]; 2) Presence of multivalent chromatin marks has been established in the unipotent and homogenous cell types[29-32]. Through combination of in-vivo and in-situ approaches, multi-valency arising in single cell has been demarcated from the one arising due to heterogeneity of the cells[33]. These studies established the presence of atypical H3K4me3/K27me3 bivalency at single cell and single allele level. It is also argued that active chromatin inhibits H3K27me3 and therefore H3K27me3 does not coincide with H3K36me3 on the same allele[34]. By extrapolation, H3K27me3 and H3K36me3 are likely to mark different alleles in the same cell across significant number of cells, if not all. 3) Further, to address the issue of heterogeneity, we hypothesize that if H3K36me3 and H3K27me3 marks are present at different levels in different cells, the genes showing variation in these marks (i.e., inferred MAE genes in the context) should also show greater cell-to-cell variation in their expression levels as compared to genes that exhibited less variation in these chromatin marks (i.e. inferred BAE genes in the context). Indeed, we tested the stochastic variation in gene expression through analysis of single cell RNA-Seq data of hLCL. We calculated the normalized expression noise of the genes (Materials and Methods) and compared that of MAE and BAE genes of comparable expression levels. As shown in the figure S11, expression noise of MAE genes was not greater than that of BAE genes. On the contrary, MAE genes exhibited lesser expression noise than the BAE genes, which also adhered to the hypothesis that MAE genes are likely to be dosage sensitive and should exhibit lesser fluctuations in their expression. Therefore, the presumed cellular heterogeneity might not entirely account for the simultaneous presence of active and inactive marks in gene-bodies of MAE genes. 5) Some of the key observations in the present study has been established on experimentally identified MAE and BAE genes through array and RNA‐seq methods. The results shown in the figure 1, and 4 can be considered very robust owing to their consistency across different experimental and inferred datasets. The results in the figure 2 could not be reproduced using experimental datasets due to lack of matching chromatin interaction datasets. However, we emphasize that these results were reproducible across different comparisons between cell-lines (figure 2 & S4). Results in figure 3 were restricted only to inferred genes due to coverage issues. The orthologous MAE/BAE genes that could be mapped near to insulator sites were insufficient[35] to carry out reliable statistical analysis. We attempted to reproduce the observations in the Figure 5 using experimentally identified MAE/BAE gene. Due to smaller sample size, data for all the chromatin states could not be obtained. However, importantly, the trend with the experimentally identified MAE/BAE genes was very similar to the ones claimed in the figure 5 (Figure S12).

Taken together, our analysis highlighted the significant dosage dependency of mitotically stable mono-allelically expressed genes, as compared to that of biallelically expressed genes, on the insulator function of CTCF protein. The correlation between evolutionary gain and loss of CTCF insulator sites with the gain and loss of mono-allelic expression together with the observation that the MAE genes are more dosage sensitive suggested that maintenance of mono-allelic expression of genes might have served as one of the potent evolutionary constraints that have shaped linearly and spatially compartmentalized genome organization. Availability of data on allelically biased expression of genes in other mammalian species would allow to infer the ancestral status of allelic expression in future, which would endow greater evolutionary insights to the phenomenon.

## Materials and Methods

### Datasets

Numbers and source of experimentally identified MAE and BAE genes and the ones inferred based on equal enrichment of H3K36me3 and H3K27me3 marks in the gene body in human LCLs, mouse LCLs and mouse ESCs are given in the Table 1. Chromatin insulator and other ChromHMM annotations for hLCLs and mESCs were taken from Ernst et al[36] and Yue et al[37] respectively. Histone modification data was obtained from ENCODE[38]. CTCF ChIA-PET datasets for hLCLs and mESCs were obtained from Tang et al[20] and Handoko et al[39] respectively. Haplotype resolved HiC dataset was obtained from Rao et al[24]. CTCF depletion datasets for mLCLs and mESCs were obtained from GSE98507 (unpublished) and Nora et al[26]. Dosage sensitive genes and CNV data were obtained from ClinGen resource (<u>https://www.ncbi.nlm.nih.gov/projects/dbvar/clingen/</u>) and Database of Genomic Variants (<u>http://dgv.tcag.ca/dgv/app/home</u>) respectively. Data for perturbation experiments were obtained from Gene Perturbation Atlas (<u>http://biocc.hrbmu.edu.cn/GPA/</u>). Single cell RNA-seq data of hLCL was obtained from GSE44618.

### Linear clustering of genes

Observed numbers of MAE-MAE, BAE-BAE, and mixed (MAE-BAE and BAE-MAE) pairs were divided by corresponding expected numbers. Expected numbers were calculated as 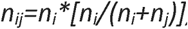, where i and j represented the allelic status of genes in a pair. Since inferred list of MAE and BAE genes had sufficient coverage of genome, we only used inferred MAE/BAE genes for this analysis. Further, hg19 and mm10 genomes were binned into 100kb windows. Density of MAE and BAE genes was calculated in those bins. Density was smoothed using ‘running.mean’ function with binwidth=3 from the “igraph” package and a linear regression line was called using ‘lm’ function in R. Bins exhibiting gene density above the linear regression line were defined as gene-clusters. Bins having only MAE (or BAE) were called as MAE (or BAE) bins. Bins with both MAE and BAE genes were called as 'mixed’ bins. For re-sampling, 9332 and 10012 genes were picked randomly and termed as MAE and BAE genes respectively. The above procedure was followed to classify bins as MAE, BAE and mixed bins. P-value was calculated using Chi-squared test for original and randomized dataset. Randomized dataset was compiled by randomly re-assigning MAE or BAE status of the genes while keeping the total number of MAE and BAE genes same as in original set.

### Kronecker delta calculation for CTCF insulation

Five genes upstream and five genes downstream were mapped around each barrier CTCF site. Kronecker delta was calculated for the pair of consecutive genes moving from upstream to downstream direction. If two genes were having same allelic status, value “1” was given. If one was MAE and other was BAE, value “0” was given. It was done for all insulator CTCF sites. Then the values were scaled from 0 to 1 and the average value of Kronecker delta was plotted.

### CTCF gap-loop-gap analysis

CTCF ChIA-PET loops which were having at least three PET pairs and spanned up to 1mb, were taken as per recommendations from Fullwood et al[40]and Li et al[19]. CTCF ChIA-PET loops that overlapped with insulator CTCF sites were taken for the analysis. Relative enrichment of TSS sites of MAE and BAE genes in the loops and the flanking regions, of same length as that of loops, was calculated. Average aggregation values for all loops were normalized with total gene count accordingly. These final values were then scaled from 0 to 1 and average values were plotted.

### Analysis of conserved and variable allelic status of genes

To compare the allelic status between cell-lines, we obtained the insulator sites for K562 cell-line and compared with that of hLCL. MAE and BAE genes that maintained their allelic status in both cell-lines and the ones that switched from mono-to‐biallelic expression and vice-versa in two cell-lines were assessed for the presence of insulator binding in the proximity (<20kb). For human-mouse comparison, orthologous gene information was taken from Ensembl. Orthologous insulator CTCF sites of human (hg19) in mouse (mm10) were obtained using UCSC liftover (with minimum ratio of bases that must remap as 0.1). Proximal presence of insulator sequence was assessed for MAE and BAE genes that maintained their allelic status in both human and mouse LCL cell-line and the ones that switched from mono-to-bi‐ allelic expression and vice-versa in two species (<20kb). P-values were calculated using Chi-squared test.

### Analysis of allele-specific chromatin loops

MAE and BAE genes from human LCL cell-line were assessed for the difference in the chromatin interaction frequency of insulator occupied sites. Due to the lack of public availability of haplotype-resolved CTCF ChIA-PET data, we overlaid CTCF ChIA-PET loops for GM12878 from Tang et al[20] onto haplotype-resolved Hi-C data for GM12878 (resolution: 5kb, normalization: VC) from Rao et al[24]. These CTCF ChIA‐PET loops were having at least three PET pairs and were up to 1Mb in length[20]. Chromatin loops with only MAE's and only BAE's gene TSSs were classified as MAE and BAE loops respectively. Squared difference between maternal and paternal loci was calculated and then divided by the maximum of their interaction frequencies for normalization. P-value was calculated using Mann Whitney U test. Since allele‐specific expression data for LCL cell-line was not available in Rao et al's article, we used RNAPII ChIA-PET data[20] to infer active and inactive alleles in Rao et al's HiC data[24]. RNA-pol2 ChIA-PET loops for GM12878 were overlaid onto maternal and paternal Hi-C datasets. Maternal-to-paternal ratio of HiC interaction frequencies for RNAPII ChIA-PET loops was calculated for each TSS site. The upper and lower quartiles of the ratio were then taken as maternally and paternally biased genes. For these M-biased and P-biased genes, allele-specific interaction frequencies of CTCF mediated loops were obtained from the HiC data. Maternal-to-paternal ratio of interaction frequencies of CTCF mediated loops was calculated and viewed as boxplots. P-value was calculated using Mann-Whitney U test.

### CTCF depletion analysis

Hi-C data (.cool format, resolution: 20kb, mm9, untreated and 2 days auxin treated) were taken from Nora et al., 2017[26]. Dip prominence scores wer used as provided by the authors. Higher dip prominence signified highly insulating boundaries. Dip prominence was mapped to 20Kb to the TSSs of MAE and BAE genes. Hi-C data (.cool format) was extracted using Cooler (<u>https://github.com/mirnylab/cooler</u>). ‘Heatmap’ function of R-package was used to plot TAD domains. To analyse the transcriptome, SRA files for LPS induced CTCF depleted B-cells were downloaded from GSE98507 and were converted into fastq format using fastq-dump (SRA Toolkit). Pilot-run fastq files were not used for further analysis. Fastq files were mapped onto mouse reference genome (mm10) and calculated differential gene expression between control and CTCF depleted samples using tophat and cufflinks without new gene/transcript discovery[41]. For mESC, RPKM values for control (untreated) and 2 days auxin-treated CTCF depleted cells were downloaded from GSE98671 [26]. Fold change was calculated with respect to control cells. Distributions of fold change (FC) of up-regulated (log2 FC > 0) and down-regulated (log2 FC < 0) MAE and BAE genes were plotted as boxplots. P-values were calculated by Mann-Whitney U test. For the analysis of expression in the neighbouring domains MAE genes with at-least 1.5 fold up‐ and down-regulation were used The nearest neighbour was taken on both sides and their RPKM values before CTCF depletion were plotted as boxplots. For Nipbl depletion analysis, we obtained the gene expression data for Nipbl and TAM‐ control from GSE93431.

### Transcriptional perturbation analysis

GPA Enrichment Tool (<u>http://biocc.hrbmu.edu.cn/GPA/analysisTool</u>) was used to calculate the proportion of MAE and BAE genes significantly perturbed (FDR<0.01) across different perturbation experiments.

### Expression noise calculation

Expression noise was calculated as coefficient of variation (standard_deviation/mean) for all the genes across 62 single human lymphoblastoid cells. The noise was then normalized against the average expression levels of genes using loess regression.

## Acknowledgement

Authors acknowledge the financial support from SERB/DST (EMR/2015/001681).

## Supplementary Figure Legends

**Figure S1.** (a) Pie charts showing the distribution of clusters with only MAE or BAE and mixed (MAE/BAE) genes in different cell-lines. hLCL: human lymphoblastic cell‐ line, mLCL: mouse lymphoblastic cell-line, mESC: mouse embryonic stem cells. P‐ value (p<2.2.e-16) was calculated using Chi-squared test for observed counts and expected counts. (b) Pie chart showing the percentage of bins with only MMAE, only PMAE and both mixed within gene-clusters. (c) Scaled average aggregation plot of Kronecker delta function for MMAE and PMAE genes.

**Figure S2. (a)** Sampled MAE and BAE genes with insignificant difference in their expression levels. **(b)** Normalized enrichment of sampled MAE and BAE genes of similar expression levels inside and around chromatin loops mediated by insulator CTCF sites in hLCL.

**Figure S3.** Normalized enrichment of MAE and BAE genes inside and around chromatin loops mediated by enhancer-linker CTCF sites in hLCL.

**Figure S4.** (a) Enrichment of MAE and BAE genes that maintained their allelic status in hLCL and HMEC cell-lines, around insulator CTCF site (within 20kb). (b) Enrichment of MAE and BAE genes that switched their allelic status in hLCL and HMEC cell-lines, around insulator CTCF site (within 20kb). P-values were calculated using Mann‐ Whitney U test. (c) Examples illustrating the switch in allelic status of genes between two cell-lines. Shown are the chromHMM tracks around: i) MYB gene, which was biallelic in hLCL but had mono-allelic expression in HMEC cell-line; and ii) NRP2 gene that had mono-allelic expression in hLCL but was expressed bi-allelically in HMEC. Solid and dashed green arcs are experimentally identified RNAPII ChIA-PET and CTCF ChIA-PET loops in hLCL respectively. Chromatin loops for HMEC were not available.

**Figure S5.** Impact of Cohesin unloading on expression of MAE and BAE genes in mESCs. Shown are the fold change in gene expression (ΔNipbl vs. Tamoxifen-control) of MAE and BAE genes.

**Figure S6.** Ratio of up-regulated to down-regulated genes after CTCF depletion in mLCL and mESC. P-values were calculated by Chi-squared test.

**Figure S7.** Proportion of MAE and BAE genes significantly (FDR<0.01) perturbed in mESCs across several different perturbations. Perturbation data was taken from Gene Perturbation Atlas (GPA)

**Figure S8.** (a) Density plot of log10 normalized squared difference in the CTCF loop interaction frequencies between the maternal and the paternal loci of MAE (Orange) and BAE (Blue) genes. (b) Box-plot of the maternal-to-paternal ratio of CTCF loop interaction frequencies of maternally and paternally biased MAE genes. Dashed line is the median of all MAE genes. All the p-values were calculated by Mann-Whitney U test (one tail).

**Figure S9.** Proportion of MAE and BAE genes overlapping with (a) dosage sensitive genes and (b) common CNVs, as annotated by ClinGen resource. P-values were calculated using Chi-squared test.

**Figure S10.** Normalized enrichment of H3K27me3 before and after CTCF depletion in the chromatin loops (and flanking regions) enclosing (a) up-regulated and (b) down‐ regulated genes.

**Figure S11.** Distribution of normalized expression noise of MAE and BAE genes of comparable expression levels (the same sampled dataset as in figure S2)

**Figure S12.** Same as in the figure 5a-b in the main text, but for the experimentally identified MAE and BAE genes in mESC.

## References

1. Gimelbrant A, Hutchinson JN, Thompson BR, Chess A: Widespread monoallelic expression on human autosomes. Science 2007, 318:1136-1140.

2. Pernis B, Chiappino G, Kelus AS, Gell PG: Cellular localization of immunoglobulins with different allotypic specificities in rabbit lymphoid tissues. J Exp Med 1965, 122:853-876.

3. Hollander GA, Zuklys S, Morel C, Mizoguchi E, Mobisson K, Simpson S, Terhorst C, Wishart W, Golan DE, Bhan AK, Burakoff SJ: Monoallelic expression of the interleukin-2 locus. Science 1998, 279:2118-2121.

4. Rhoades KL, Singh N, Simon I, Glidden B, Cedar H, Chess A: Allele-specific expression patterns of interleukin-2 and Pax-5 revealed by a sensitive single-cell RT-PCR analysis. Curr Biol 2000, 10:789-792.

5. Brady BL, Steinel NC, Bassing CH: Antigen receptor allelic exclusion: an update and reappraisal. J Immunol 2010, 185:3801-3808.

6. Chess A, Simon I, Cedar H, Axel R: Allelic inactivation regulates olfactory receptor gene expression. Cell 1994, 78:823-834.

7. Chess A: Mechanisms and consequences of widespread random monoallelic expression. Nat Rev Genet 2012, 13:421-428.

8. Lewcock JW, Reed RR: A feedback mechanism regulates monoallelic odorant receptor expression. Proc Natl Acad Sci U S A 2004, 101:1069-1074.

9. Saleh A, Davies GE, Pascal V, Wright PW, Hodge DL, Cho EH, Lockett SJ, Abshari M, Anderson SK: Identification of probabilistic transcriptional switches in the Ly49 gene cluster: a eukaryotic mechanism for selective gene activation. Immunity 2004, 21:55-66.

10. Meaburn EL, Schalkwyk LC, Mill J: Allele-specific methylation in the human genome: implications for genetic studies of complex disease. Epigenetics 2010, 5:578-582.

11. Shoemaker R, Deng J, Wang W, Zhang K: Allele-specific methylation is prevalent and is contributed by CpG-SNPs in the human genome. Genome Res 2010, 20:883-889.

12. Mancini-Dinardo D, Steele SJ, Levorse JM, Ingram RS, Tilghman SM: Elongation of the Kcnq1ot1 transcript is required for genomic imprinting of neighboring genes. Genes Dev 2006, 20:1268-1282.

13. Sleutels F, Zwart R, Barlow DP: The non-coding Air RNA is required for silencing autosomal imprinted genes. Nature 2002, 415:810-813.

14. Takizawa T, Gudla PR, Guo L, Lockett S, Misteli T: Allele-specific nuclear positioning of the monoallelically expressed astrocyte marker GFAP. Genes Dev 2008, 22:489-498.

15. Armelin-Correa LM, Gutiyama LM, Brandt DY, Malnic B: Nuclear compartmentalization of odorant receptor genes. Proc Natl Acad Sci U S A 2014, 111:2782-2787.

16. Nag A, Savova V, Fung HL, Miron A, Yuan GC, Zhang K, Gimelbrant AA: Chromatin signature of widespread monoallelic expression. Elife 2013, 2:e01256.

17. Edwards CA, Ferguson-Smith AC: Mechanisms regulating imprinted genes in clusters. Curr Opin Cell Biol 2007, 19:281-289.

18. Kieffer-Kwon KR, Tang Z, Mathe E, Qian J, Sung MH, Li G, Resch W, Baek S, Pruett N, Grontved L, et al: Interactome maps of mouse gene regulatory domains reveal basic principles of transcriptional regulation. Cell 2013, 155:1507-1520.

19. Li G, Ruan X, Auerbach RK, Sandhu KS, Zheng M, Wang P, Poh HM, Goh Y, Lim J, Zhang J, et al: Extensive promoter-centered chromatin interactions provide a topological basis for transcription regulation. Cell 2012, 148:84-98.

20. Tang Z, Luo OJ, Li X, Zheng M, Zhu JJ, Szalaj P, Trzaskoma P, Magalska A, Wlodarczyk J, Ruszczycki B, et al: CTCF-Mediated Human 3D Genome Architecture Reveals Chromatin Topology for Transcription. Cell 2015, 163:1611-1627.

21. Zhang Y, Wong CH, Birnbaum RY, Li G, Favaro R, Ngan CY, Lim J, Tai E, Poh HM, Wong E, et al: Chromatin connectivity maps reveal dynamic promoter‐enhancer long-range associations. Nature 2013, 504:306-310.

22. Gotea V, Visel A, Westlund JM, Nobrega MA, Pennacchio LA, Ovcharenko I: Homotypic clusters of transcription factor binding sites are a key component of human promoters and enhancers. Genome Res 2010, 20:565-577.

23. Kurukuti S, Tiwari VK, Tavoosidana G, Pugacheva E, Murrell A, Zhao Z, Lobanenkov V, Reik W, Ohlsson R: CTCF binding at the H19 imprinting control region mediates maternally inherited higher-order chromatin conformation to restrict enhancer access to Igf2. Proc Natl Acad Sci U S A 2006, 103:10684-10689.

24. Rao SS, Huntley MH, Durand NC, Stamenova EK, Bochkov ID, Robinson JT, Sanborn AL, Machol I, Omer AD, Lander ES, Aiden EL: A 3D map of the human genome at kilobase resolution reveals principles of chromatin looping. Cell 2014, 159:1665-1680.

25. Lin S, Ferguson-Smith AC, Schultz RM, Bartolomei MS: Nonallelic transcriptional roles of CTCF and cohesins at imprinted loci. Mol Cell Biol 2011, 31:3094-3104.

26. Nora EP, Goloborodko A, Valton AL, Gibcus JH, Uebersohn A, Abdennur N, Dekker J, Mirny LA, Bruneau BG: Targeted Degradation of CTCF Decouples Local Insulation of Chromosome Domains from Genomic Compartmentalization. Cell 2017, 169:930-944 e922.

27. Toyooka Y, Shimosato D, Murakami K, Takahashi K, Niwa H: Identification and characterization of subpopulations in undifferentiated ES cell culture. Development 2008, 135:909-918.

28. Hayashi K, de Sousa Lopes SMC, Tang F, Lao K, Surani MA: Dynamic equilibrium and heterogeneity of mouse pluripotent stem cells with distinct functional and epigenetic states. Cell Stem Cell 2008, 3:391-401.

29. Dahl JA, Reiner AH, Klungland A, Wakayama T, Collas P: Histone H3 lysine 27 methylation asymmetry on developmentally-regulated promoters distinguish the first two lineages in mouse preimplantation embryos. PLoS One 2010, 5:e9150.

30. Roh TY, Cuddapah S, Cui K, Zhao K: The genomic landscape of histone modifications in human T cells. Proc Natl Acad Sci U S A 2006, 103:15782-15787.

31. Alder O, Lavial F, Helness A, Brookes E, Pinho S, Chandrashekran A, Arnaud P, Pombo A, O’Neill L, Azuara V: Ring1B and Suv39h1 delineate distinct chromatin states at bivalent genes during early mouse lineage commitment. Development 2010, 137:2483-2492.

32. Vastenhouw NL, Zhang Y, Woods IG, Imam F, Regev A, Liu XS, Rinn J, Schier AF: Chromatin signature of embryonic pluripotency is established during genome activation. Nature 2010, 464:922-926.

33. Brookes E, de Santiago I, Hebenstreit D, Morris KJ, Carroll T, Xie SQ, Stock JK, Heidemann M, Eick D, Nozaki N, et al: Polycomb associates genome-wide with a specific RNA polymerase II variant, and regulates metabolic genes in ESCs. Cell Stem Cell 2012, 10:157-170.

34. Schmitges FW, Prusty AB, Faty M, Stutzer A, Lingaraju GM, Aiwazian J, Sack R, Hess D, Li L, Zhou S, et al: Histone methylation by PRC2 is inhibited by active chromatin marks. Mol Cell 2011, 42:330-341.

35. Zwemer LM, Zak A, Thompson BR, Kirby A, Daly MJ, Chess A, Gimelbrant AA: Autosomal monoallelic expression in the mouse. Genome Biol 2012, 13:R10.

36. Ernst J, Kheradpour P, Mikkelsen TS, Shoresh N, Ward LD, Epstein CB, Zhang X, Wang L, Issner R, Coyne M, et al: Mapping and analysis of chromatin state dynamics in nine human cell types. Nature 2011, 473:43-49.

37. Yue F, Cheng Y, Breschi A, Vierstra J, Wu W, Ryba T, Sandstrom R, Ma Z, Davis C, Pope BD, et al: A comparative encyclopedia of DNA elements in the mouse genome. Nature 2014, 515:355-364.

38. Consortium EP: An integrated encyclopedia of DNA elements in the human genome. Nature 2012, 489:57-74.

39. Handoko L, Xu H, Li G, Ngan CY, Chew E, Schnapp M, Lee CW, Ye C, Ping JL, Mulawadi F, et al: CTCF-mediated functional chromatin interactome in pluripotent cells. Nat Genet 2011, 43:630-638.

40. Fullwood MJ, Liu MH, Pan YF, Liu J, Xu H, Mohamed YB, Orlov YL, Velkov S, Ho A, Mei PH, et al: An oestrogen-receptor-alpha-bound human chromatin interactome. Nature 2009, 462:58-64.

41. Trapnell C, Roberts A, Goff L, Pertea G, Kim D, Kelley DR, Pimentel H, Salzberg SL, Rinn JL, Pachter L: Differential gene and transcript expression analysis of RNA-seq experiments with TopHat and Cufflinks. Nat Protoc 2012, 7:562-578.

42. Gendrel AV, Attia M, Chen CJ, Diabangouaya P, Servant N, Barillot E, Heard E: Developmental dynamics and disease potential of random monoallelic gene expression. Dev Cell 2014, 28:366-380.

43. Nag A, Vigneau S, Savova V, Zwemer LM, Gimelbrant AA: Chromatin Signature Identifies Monoallelic Gene Expression Across Mammalian Cell Types. G3 (Bethesda) 2015, 5:1713-1720.

44. Rozowsky J, Abyzov A, Wang J, Alves P, Raha D, Harmanci A, Leng J, Bjornson R, Kong Y, Kitabayashi N, et al: AlleleSeq: analysis of allele-specific expression and binding in a network framework. Mol Syst Biol 2011, 7:522.

